# Trigeminal ganglion and tooth innervation modifications following genetic and pharmacological Nogo-A inhibition

**DOI:** 10.1101/2024.06.04.597304

**Authors:** Laurence Pirenne, Anamaria Balic, Ilaria De Santis, Alessandro Bevilacqua, Chai Foong Lai, Pierfrancesco Pagella, Martin E. Schwab, Thimios A. Mitsiadis

**Affiliations:** Institute of Oral Biology, Centre of Dental Medicine, Faculty of Medicine, University of Zurich, Zurich, Switzerland; Department of Computer Science and Engineering, University of Bologna, Bologna, Italy; Brain Research Institute, University of Zurich, Zurich, Switzerland; Institute for Regenerative Medicine, University of Zurich, Zurich, Switzerland

## Abstract

Nogo-A is a major regulator of neural development and regeneration, but its role in tooth innervation remains largely unknown. Neurons from trigeminal ganglia support teeth homeostasis and regeneration, and disorders of their function could have significant pathophysiological consequences. In this study, we show that Nogo-A is expressed in the trigeminal ganglia and in the neurons innervating the teeth, and that its deletion affects both the number and patterning of neurons in teeth. In organotypic cultures, Nogo-A blocking antibodies affect the trigeminal ganglia-derived neuronal outgrowths and allow premature innervation of tooth germs. RNA sequencing analysis revealed that Nogo-A deletion induces alterations linked to functions at synapses and interference with neurotrophin signalling during the differentiation and maturation of trigeminal neurons. Taken together, these results reveal for the first time the importance of Nogo-A as a major regulator of tooth innervation and point to its potential as a clinical therapeutic target.

## Introduction

Nogo-A is a 130-kDa transmembrane protein encoded by the *Reticulon-4 (Rtn4)* gene (1). Alternative splicing of *Rtn4* gives rise to the Nogo-B (40kDa) and Nogo-C (20kDa) isoforms (1, 2). All three isoforms share a conserved C-terminal domain but have different N-terminal domains (1, 2). Nogo-A is located in the endoplasmic reticulum membrane, as well as on the cell surface (1, 3). At the cell surface, Nogo-A acts as a ligand for several multi-subunit receptor complexes and triggers signalling cascades (4, 5). Nogo-66 (located in the C- terminal part of all Nogo isoforms) and Δ20 (found only in the N-terminal part of the Nogo-A isoform) are the two main functional domains of the protein (3, 6). The Δ20 domain interacts with S1PR2 (Sphingosine-1-Phosphate Receptor 2) (4), whereas Nogo-66 interacts with NgR1 (Nogo Receptor 1) (7). Often, NgR1 forms complexes with other transmembrane receptors, including LINGO-1 (immunoglobulin-like domain-containing protein-1) and p75^NTR^ (p75 neurotrophin receptor) (8, 9). Nogo-A binding to its various receptor complexes activates RhoA and ROCK (Rho-Associated Kinase) signalling that promotes the disorganisation of the actin cytoskeleton and growth cone collapse (9, 10). Nogo-A signalling also leads to transcriptional changes and the suppression of growth-related genes (11–13).

Nogo-A is an important regulator of neurite growth in the central nervous system (CNS) (14, 15). During development, Nogo-A expression in neurons is involved in their migration (16), differentiation (17), outgrowth, and branching (18, 19). In the adult brain, Nogo-A is predominantly expressed by oligodendrocytes and stabilises the existing neuronal networks (12, 20). Upon CNS injury, Nogo-A inhibits the regeneration of axons; in animal models, its genetic ablation (i.e., Nogo-A knockout mice) or inactivation by antibodies and blocking peptides enhances regenerative axonal growth and functional recovery after spinal cord or brain injuries (5). Indeed, Nogo-A is a specific target for the treatment of CNS lesions in humans in ongoing clinical trials; https://nisci-2020.eu. Although the regenerative potential of Nogo-A neutralization could also be beneficial for peripheral highly innervated tissues and organs, the role and functions of Nogo-A outside the CNS remain largely unknown and unexplored.

Teeth are richly innervated organs, and numerous studies have been conducted to understand the role of neurons in the homeostasis, pathology, and regeneration of the dental pulp and periodontal tissues (21–24). The tooth-specific neuronal network is composed of both sensory and sympathetic nerve fibres originating from the trigeminal ganglion (tgg) and superior cervical ganglion, respectively (22, 25). The sensory fibres respond to electric, thermal, mechanical, and chemical stimuli by providing tooth pain sensation (22), while the sympathetic innervation regulates the vasculature and controls blood flow within dental tissues (26). Tooth innervation is also important for immunomodulation (27–30) and stem cell behaviour (31, 32) in pathological conditions, as well as during dental tissue repair and regeneration processes. Tooth-specific pathologies, including traumatic injuries, carious decays, and periodontal diseases, lead to increased neuronal sensitivity and elevated pain threshold (33, 34).

Innervation occurs concomitantly with organogenesis (35) and is often critical for the development of organs such as taste buds (36), cochlea (37), and salivary glands (38). In contrast to other organs, the innervation of the developing tooth germs is delayed and starts at advanced morphogenetic stages (i.e., the late bell stage), as nerve fibres invade the dental pulp after initial dentine deposition (39–41). These neurons rapidly expand and form networks within the dental pulp of the developing teeth (42, 43). In mature teeth, the nerve fibres form a dense neuronal plexus at the periphery of the pulp, in close proximity to the dentine-secreting odontoblasts (44). Previous studies have revealed the importance of specific signalling molecules in tooth innervation, including members of the neurotrophin and semaphorin families. Neurotrophins, including nerve growth factor (NGF), brain-derived neurotrophic factor (BDNF), neurotrophin-3 (NT3), and neurotrophin-4/5 (NT4/5), are factors essential for the development, differentiation, sprouting, and survival of the peripheral sensory and sympathetic neurons (45). The neuro-attractive cues of neurotrophins serve to guide the nerve fibres towards their targets, including the dental tissues (46–48). In contrast, members of the semaphorin family (49), such as semaphorin-3A (Sema3A), are neurite repulsive molecules that inhibit the innervation of the dental pulp in the developing tooth germs (50, 51). The neuroregulatory properties of these molecules allowed the development of innovative therapeutic strategies for the regeneration of dental tissues (52, 53).

Since Nogo-A acts as a neuron repulsive signal, it is likely that it is also involved in the development and establishment of tooth innervation. In this study, we analysed the distribution patterns of Nogo-A in embryonic and postnatal tgg, as well as in the developing and erupted teeth of rodents, using immunostaining and state-of-the-art imaging techniques. We also assessed the effects of Nogo-A on tooth innervation using Nogo-A knockout (KO) mice and antibodies in organotypic explant cultures. Finally, we performed transcriptomic analyses to examine the effects of Nogo-A deletion on the molecular signature of the tgg. The present results identify Nogo-A as an important regulator of tooth innervation and indicate its potential as a future clinical therapeutic tool for peripheral innervated organs.

## Materials and Methods

### Animals and ethics statement

Animal housing and experimentation were performed according to the Swiss Animal Welfare Law and in compliance with the regulations of the Cantonal Veterinary Office of Zurich (License Number 203/2020). Wild type C57/BL6J, Nogo-A KO (54) (kindly provided by Prof. Schwab), and ROSA26-mT/mG (R26-mTmG, JAX stock #007676) (55) mice were used at different embryonic and postnatal stages, ranging from E13.5 to E15.5 and P4 to P25. The day of detection of the vaginal plug was marked as embryonic (E) day 0.5, and the day of birth was considered as postnatal (P) day 0. The embryos were further staged according to limb and craniofacial morphological criteria.

### Tissue collection

#### Histological analyses

Whole embryos were collected in cold phosphate-buffered saline (PBS) and fixed overnight in 4% paraformaldehyde (PFA; Sigma-Aldrich, Buchs, Switzerland, #16005). The following day, embryos were washed with PBS and incubated in 30% sucrose until sinking. The embryos were then frozen embedded in Tissue-Tek^®^ O.C.T. compound (OCT) (Sakura Finetek, Alphen aan den Rijn, The Netherlands, #4583). Postnatal tgg and first lower molars were dissected in cold PBS under a M80 Leica stereomicroscope. Ganglia were directly frozen embedded in OCT, while the molars were first fixed in 4% PFA overnight at 4°C, and then decalcified in 0.5M ethylenediaminetetraacetic acid (EDTA; NeoFroxx, Einhausen, Germany, #LC-10442.5) for 2-3 weeks. Decalcified molars were washed in PBS and then processed for frozen embedding in OCT as described for embryos, or gradually dehydrated in increasing concentrations of ethanol, followed by clearing in xylol and paraffin embedding.

OCT samples were cut at either 10µm (thin sections) or at 40µm (thick sections). Paraffin samples were cut at 7µm.

#### Whole mount immunostaining

Dental pulps were isolated from WT and Nogo-A KO mice at P4, P7 and P25. The coronal portion of the dental pulp was isolated from the first lower molars of P4 and P7 pups and fixed in 4% PFA for 15 minutes. The whole first lower molars of P25 mice were extracted in cold PBS, fixed in 4% PFA (Sigma-Aldrich, #16005) for 15 minutes and then decalcified in 0.5M EDTA for a week. Once the hard tissues had softened enough, the whole intact dental pulps were manually dissected out of it.

### Immunohistochemistry

Immunoperoxidase staining on thin cryosections and paraffin sections was performed as previously described (46). Cryosections were incubated in PBS and paraffin sections were deparaffinised and rehydrated before being exposed to a 0.5% H_2_O_2_ (Sigma-Aldrich, #216763) in methanol. Sections were then processed for antigen retrieval in boiling TE-9 followed by permeabilization in 1% triton in PBS. Next, sections were blocked with 1% BSA (Carl Roth, Arlesheim, Switzerland, #0163.2) and 10% goat serum (ThermoFisher Scientific, Zürich, Switzerland, #16210064) in PBS for 2 hours at room temperature, followed by the overnight incubation with anti-Nogo-A (rabbit, see **Table 1**) at 4°C in a humid atmosphere. Primary antibody labelling was detected and visualized by VECTASTAIN® ABC-HRP Kit, peroxidase rabbit IgG (Vector Laboratories, Zürich, Switzerland, #PK-4001) and the AEC substrate kit (Vector Laboratories, #SK4200), that were used following manufacturer’s instructions. After staining, sections were counterstained with haematoxylin (Morphisto, Muttenz, Switzerland, #18417) and mounted with glycergel mounting medium (Agilent Dako, Basel, Switzerland, #C0563).

**Table 1:**
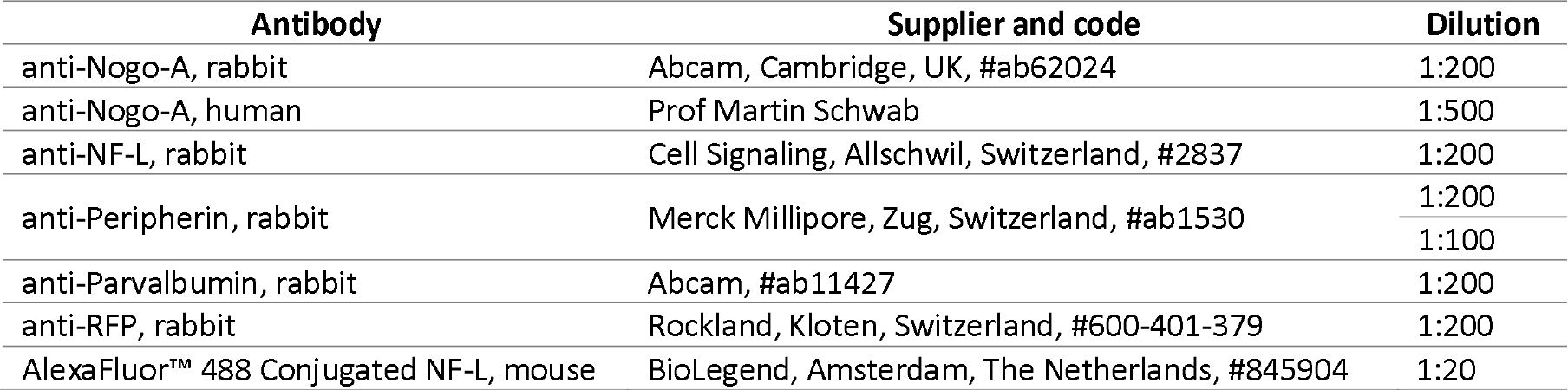
List of primary antibodies used in this study, with the associated information on the supplier and the dilution.

### Immunofluorescent staining

Immunofluorescent (IF) staining was performed on thin and thick cryosections, whole dental pulp tissues and co-cultured tgg / tooth-germs. <u>Thin cryosections</u> were blocked with 10% goat serum (ThermoFisher Scientific, #16210064) and 1% BSA in PBS (Carl Roth, #0163.2) for 30 minutes, and incubated overnight at 4°C with primary antibodies diluted in blocking solution. The primary antibodies used are listed in **Table 1** . Primary antibody was then washed with PBS and samples were incubated with the secondary antibodies diluted 1:500 in 1% BSA-PBS, for 1 hour at room temperature. Secondary antibodies used are listed in **Table 2** . Nuclei were stained with DAPI (1 µg/mL, Tocris Bioscience, #5748), and after extensive washing with PBS, the sections were mounted with fluorescence mounting medium (Agilent Dako, #S3023). <u>Thick cryosections</u> were processed similarly to thin cryosections, except they were blocked with 10% goat serum / 1%BSA in PBS / 0.05% saponin (Fluka, Buchs, Switzerland, #84510) overnight at 4°C, and the incubation with the secondary antibody was overnight at 4°C. <u>Whole dental pulps and cultured tgg</u> were also blocked with 10% goat serum / 1%BSA in PBS / 0.05% saponin overnight at 4°C, then incubated with the anti-peripherin antibody (see **Table 1**) for 72 hours at 4°C on a shaker. Following PBS washes, the dental pulps were incubated with the neurofilament-light AlexaFluor™ 488 conjugated antibody (see **Table 1** ), goat anti-rabbit AlexaFluor™ 568 secondary antibody (see **Table 2** ) and DAPI (1 µg/mL) for 72 hours at 4°C on a shaker. Following PBS washes, dental pulps were incubated at 4°C in HistoDenz™ (Sigma-Aldrich, #D2158) until cleared, and mounted on a glass slide moulded with silicone paste KORASILON™ (Carl Roth, #APA71). Images were acquired at the confocal microscope CLSM Leica Stellaris 5 supplemented with the LAS X software. Whole dental pulps were imaged as z-stacks with a 4.28µm z-step and are shown as a maximum projection using Imaris 9.9 (Bitplane AG, Oxford Instruments, Switzerland). Overview images of cultured tgg were acquired with a Widefield Leica THUNDER microscope supplemented with the LAS X software. <u>Staining in microfluidic devices</u> was performed as previously described (56).

**Table 2:** List of secondary antibodies used in this study, with the associated information on the supplier and dilution at which they were used.

| Antibody | Supplier and code | Dilution |
| --- | --- | --- |
| Anti-Rabbit, Peroxidase | Vector Laboratories, Zürich, Switzerland, #PK-4001 | 1:200 |
| anti-Human AlexaFluor™ 647, goat | ThermoFisher Scientific, #A-21445 | 1:500 |
| anti-Rabbit AlexaFluor™ 488, goat | ThermoFisher Scientific, #A-27034 | 1:500 |
| anti-Rabbit AlexaFluor™ 568, goat | Invitrogen, Zürich, Switzerland #A-11036 | 1:100 to 1:500 |

### Segmentation and quantification of the neuronal network of whole dental pulp tissues

The analysis of the molar 3D neuronal network was performed as described in (57). The image processing procedures were implemented in MATLAB^®^ (R2021b v.9.11.0, The MathWorks). Briefly, the neuronal network was first segmented in 2D sections, then a 3D binary mask was created by simple juxtaposition of single 2D section binary masks and the segmented network was skeletonised. Non-skeletonized and skeletonized network masks were used to quantify network surface and volume, or network nodes and branches, respectively. The data was then plotted in Python using the Statannotations (58) and Seaborn (59) modules. In total 32 samples obtained from WT and KO mice at P4, P7 and P25 were used for analysis. Median, first (Q1) and third (Q3) quartiles, and 1.5× the interquartile range (IQR= Q3 – Q1) were determined for each measurement. All the samples (8 in total) which fell outside the range defined by (Q1 - 1.5× IQR) and (Q3 + 1.5× IQR) were excluded.

The statistical analysis was performed on total 24 samples, with at least 3 independent samples in each group. Statistical significance was assessed with a Two-sided Mann– Whitney U test (ns: p > 0.05, *: 0.01 < p ≤ 0.05, **: 0.001 < p ≤ 0.01, ***: 0.0001 < p ≤ 0.001, ****: p ≤ 0.0001).

### Organotypic cultures

#### Tgg cultures

Tgg were dissected from E13.5-E15.5 WT and Nogo-A KO embryos and cultured on glass coverslips coated with 20 µg/mL poly-D-lysine (Sigma-Aldrich, #P0296) and 2 µg/mL Laminin (Sigma-Aldrich, #L2020). Cultures were incubated at 37°C and 5% CO_2_ in Neurobasal Medium Plus (Gibco, ThermoFisher Scientific, #A3582901) supplemented with 1x B-27 (Gibco, ThermoFisher Scientific, #A3582901), 20 mM Glutamine (Gibco, ThermoFisher Scientific, #25030), 100 ng/mL NGF (PeproTech, ThermoFisher Scientific, #AF45001), 10U penicillin and 0.1 mg/mL streptomycin (Sigma-Aldrich, #P4333). To inhibit mitosis of non-neuronal cells, 0.25 pM cytosine arabinoside (Sigma-Aldrich, #C1768) was added to the media. Tgg were cultured for 2 days and fixed in 4% PFA. They were imaged with a DMIL LED Leica microscope supplemented with a DFC 7000T Leica camera and the LAS X software.

### Tgg and tooth germ co-cultures

Tgg were dissected from E14 R26-mTmG mouse embryos, while first molar tooth germs were dissected from lower and upper jaws of E14.5 WT and Nogo-A KO embryos. Glass coverslips were coated with 20 µg/mL poly-D-lysine and 2 µg/mL laminin, and then covered with low melting agarose (CondaLab, Basel, Switzerland, #8010). A hole was made in the centre of the coverslip using a 5 mm biopsy punch. In the first experimental setup (long-range cultures), one tgg and one WT tooth germ were placed together in the hole approximately 1.5 mm away from each other. In the second setup (short-range cultures), one tgg and one WT or Nogo-A KO tooth germ were placed in immediate proximity to each other. Cultures were then incubated at 37°C and 5% CO_2_ in Neurobasal Medium Plus supplemented with 1x B-27, 20 mM glutamine, 100 ng/mL NGF, 50 µg/mL of ascorbic acid (Sigma-Aldrich, #A4544), 10% foetal bovine serum (FBS; Sigma-Aldrich, #F7524), 10U penicillin, 0.1 mg/mL streptomycin, and 2.5 µg/mL fungizone. Long-range cultures were treated with either 10 µg/mL anti-Nogo-A mouse monoclonal antibody (antibody 11C7 (3), kindly provided by Prof. Schwab) or with 10 µg/mL mouse IgG1 isotype control antibody (Invitrogen, #02-6100). No treatment was added in the short-range cultures. Long-range cultures were stopped after 6 days, while short-range cultures were stopped after 4 days by fixing the tissue with warm 4% PFA for 15 minutes.

### Tgg and tooth germ co-cultures in microfluidic devices

Tgg were obtained from E13.5-15.5 WT mouse embryos, while the tooth germs were dissected from E16.5 WT and E16.5 Nogo-A KO mouse embryos. Microfluidic devices (Millipore, Buchs, Switzerland, #AX150) were punched with a 1 mm diameter biopsy punch. One punch was made in the middle on the right side of the microchannels for the tgg and 2 punches were made on the left side, one in the top for the WT tooth germ and one down for the Nogo-A KO tooth germ. Devices were assembled and coated with 0.1 mg/mL poly-D-lysine and 5 μg/mL Laminin as previously described (56). Tgg were cultured in the same culture medium as previously described (56). Tooth germs were cultured in a medium containing DMEM (high glucose 4.5 mg/mL) (Sigma, #D5796), 20% FBS, 20 mM glutamine, 50 µg/mL ascorbic acid, and 0.1 mg/mL streptomycin. Co-cultures were maintained for 10 days at 37°C and 5% CO_2_, and media were changed every 72 hours. After culture, samples were washed with PBS and fixed with 4% PFA for 15min.

### Segmentation and Sholl analysis of cultured tgg explants

The tgg neuronal network was segmented by training the LABKIT plugging of ImageJ (60) to recognise the network from bright field images or the RFP channel images. The segmentations were then imported to Python and the Sholl analysis was performed using the Skan module (61). In each experiment, four samples of each condition were analysed. The data was plotted in Python using the Seaborn module (59) as a line plot representing the mean number of intersections against the radial distance from the tgg centre. The light-coloured area represents the 95% confidence interval.

### High Throughput Sequencing (NGS) RNA sequencing

Total RNA was extracted from E14 and P25 WT and Nogo-A KO tgg, using TRIzol^TM^ reagent (Thermofischer, #15596026). Briefly, tissues were broken down using a motor pestle. Prior incubation with phenol/chloroform/iso-amyl alcohol (Sigma, #P1944), 2M sodium acetate at pH4 was added for 10 minutes at room temperature. After centrifugation, the aqueous phase was transferred and further cleaned with chloroform. RNA was then precipitated using isopropanol and resuspended in RNase free H_2_O (Sigma, #W4502). The RNA integrity number (RIN) and quantification for total RNA were determined using a 4200 TapeStation system (Agilent Technologies, Basel, Switzerland). In total 4 samples for each genotype and stage, with a RIN > 8 were selected for downstream analysis. Libraries were prepared using the TruSeq stranded mRNA library (Illumina, Zürich, Switzerland, #20020595) assay and sequenced on an illumina NovaSeq 6000 system at the Functional Genomics Centre of Zurich (FGCZ). Adapter and quality trimming of the raw reads were performed using fastp (v0.20.0) (62) considering a minimum read length of 18 nts and a quality cut-off of 20. The resulting high-quality reads were aligned to the mouse reference genome (build GRCm39 /Release_M26-2021-04-20) and gene level expression quantification was carried out using Kallisto (Version 0.46) (63). The DESeq2 package (v1.32.0) was used to obtain differentially expressed gene between WT and KO samples (64). Differential usage of exons and splice junctions was analysed using JunctionSeq (v1.21.0) (65). Differentially expressed genes and isoforms were screened based on |log_2_Ratio|>l10.5 and p.adjl1<l10.05 and processed for enrichment analysis using the EnrichR website and the BioPlanet2019 database (66).

## Results

### Trigeminal neurons express Nogo-A during embryonic development and postnatally

To investigate the expression of Nogo-A in the innervation of the developing tooth, we first performed immunostaining for Nogo-A in the tgg (**Fig. 1** ). At embryonic day 13.5 (E13.5), Nogo-A was clearly expressed in the tgg, when peripheral projections are growing towards the tooth germs of the mouse embryos (**Fig. 1A and B** ). Nogo-A staining was co-localised with the neurofilament (NF-L) staining (**Fig. 1C**) in the neuronal cell bodies of the tgg, where it was detected at varying expression levels (**Fig. 1B and C** ). Variable Nogo-A expression levels were maintained in the tgg neuronal cell bodies and the trigeminal neurons of postnatal day 25 (P25) mice (**Fig. 1D and F** ). As expected, Nogo-A immunoreactivity was absent in the tgg of Nogo-A KO mice (**Fig. 1E**).

**Figure 1:**
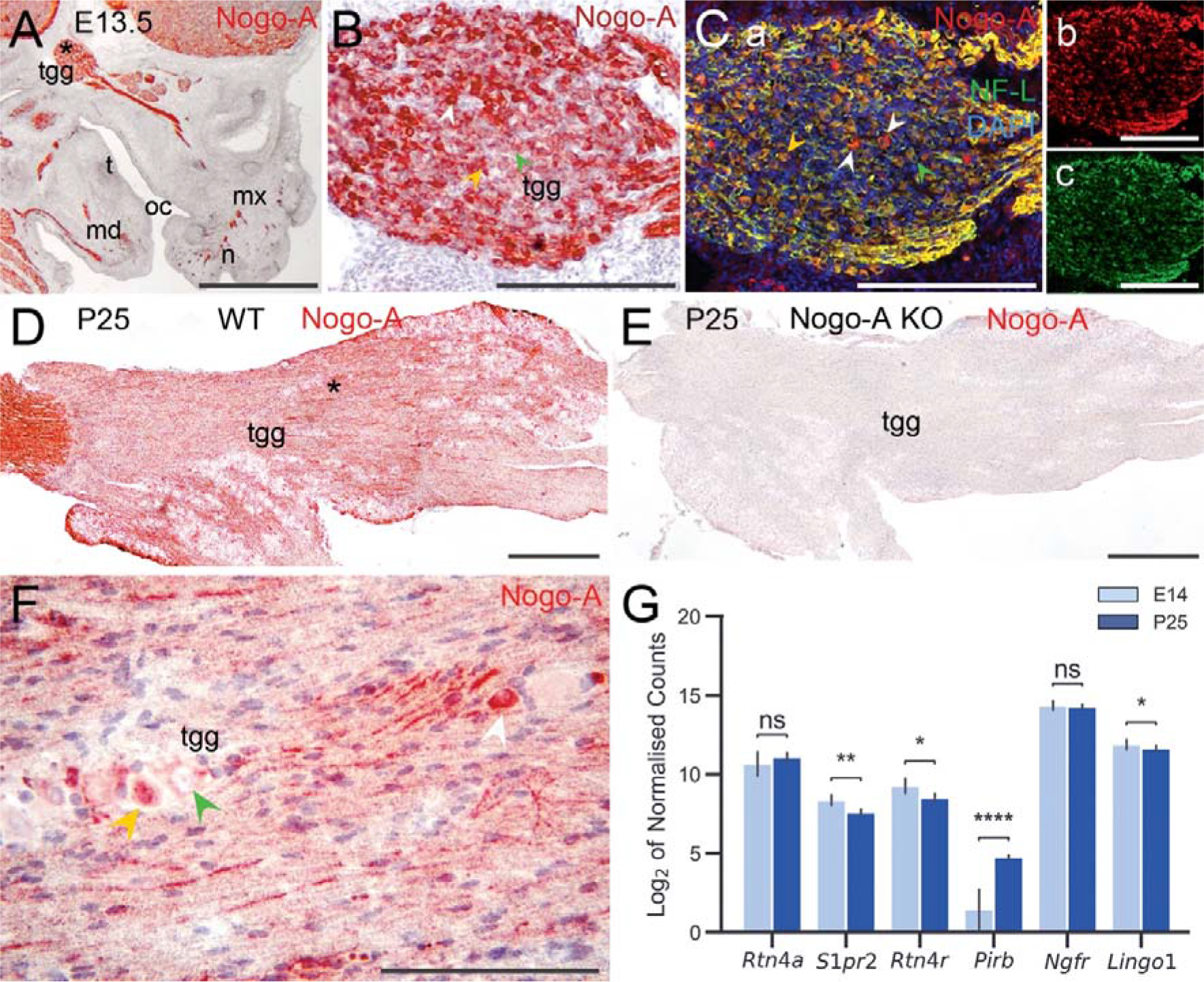
Nogo-A expression in the trigeminal ganglia of embryonic and postnatal mice. (A- B, D-F) Immunohistochemistry analysis of Nogo-A expression in the tgg of embryonic day 13.5 (E13.5) WT mouse embryos (A, B), and in WT (D, F) and Nogo-A KO (E) tgg at postnatal day 25 (P25). The asterisk in (A) indicates the area of higher magnification images in (B, C), while the asterisk in (D) indicates the area of higher magnification in (F). **(C)** Immunofluorescent co-staining of Nogo-A (red colour) and neurofilament-light chain (NF-L, green colour) showing the distribution of these two proteins at E13.5 WT tgg. Yellow colour in the merged image (Ca) indicates cells in which Nogo-A and NF-L are co-localised. White, yellow, and green arrowheads indicate high, intermediate, and low levels of Nogo-A expression, respectively. **(G)** Comparative analysis of mRNA expression levels of Nogo-A and its receptors in the trigeminal ganglion of WT mice at E14 (light blue colour) and P25 (dark blue colour). Expression levels are shown as Log_2_ of specific counts normalised to the total count of each sample. Error bars represent the standard deviation. False Discovery Rate: ns, p ≤ 0.05; *, p ≤ 0.05; **, p ≤ 0.01; ***, p ≤ 0.001. Abbreviations: md, mandibular process; mx, maxillary process; n, nose; oc, oral cavity; t, tooth germ; tgg, trigeminal ganglion. Scale bars: 1 mm in (A, D, E), 200 µm in (B, C, F).

Transcriptomics analyses further showed similar expression levels of the Nogo-A isoform (represented by *Rtn4a* expression) in the tgg of embryonic and postnatal mice (**Fig. 1G** ), which also expressed several of the Nogo-A receptors and co-receptors, albeit at different levels (**Fig. 1G** ). The expression levels of *S1pr2* and *Rtn4r* receptors, as well as *Lingo1* co-receptor, were higher in the embryonic tgg, while high expression levels of *PirB* receptor marked the adult tgg. The expression levels of *Ngfr* co-receptor were similar between the two stages. These observations corroborate previous reports on the presence of Nogo-A and NgR1 at the mRNA level in the developing rat and mouse tgg (67, 68), and indicate stage-dependent expression pattern of Nogo-A and its receptors in the tgg.

### Nogo-A is expressed in nerve fibres during embryonic and postnatal tooth development

As trigeminal neurons express Nogo-A, we sought out if Nogo-A is transported from the cell bodies to the peripheral axons innervating the tooth and performed immunostaining for Nogo-A in the developing E14.5 tooth germs and the fully erupted P25 teeth (**Fig. 2** ). Peripheral nerve fibres positive for both Nogo-A (**Fig. 2A and B** ) and NF-L (**Fig. 2B** ) were observed in the close vicinity of the tooth germs, around the condensed dental mesenchyme. While we detected Nogo-A staining in the dental epithelium (**Fig. 2A and B** ), which is in line with previous observations ((69) and our unpublished data), the majority of Nogo-A expression localized at the border with the dental mesenchyme (**Fig. 2B** ). Interestingly, a low level of Nogo-A expression was observed throughout the embryonic mandibular tissues except in the dental mesenchyme (**Fig. 2B**).

**Figure 2:**
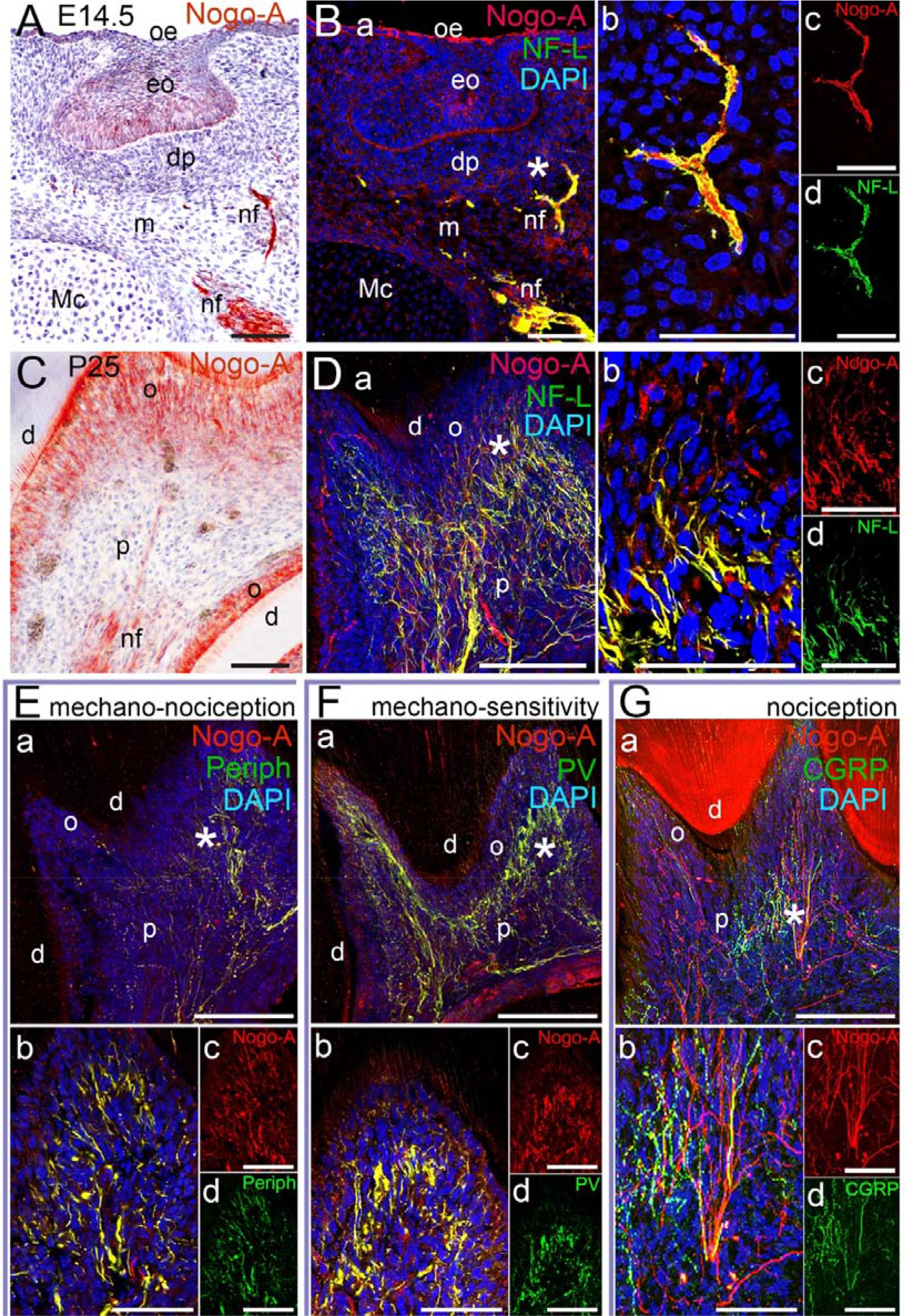
Patterns of Nogo-A expression in dental tissues of developing embryonic and postnatal mouse teeth. Immunostaining showing the distribution of the Nogo-A protein in the E14.5 WT molar tooth germs (A-B), and the P25 WT molars (C-G). **(A)** Immunohistochemistry showing the localisation of Nogo-A in the nerve fibres (nf) and the enamel organ (eo) of an E14.5 mouse tooth germ. **(B)** Immunofluorescent staining of Nogo-A (red colour in Ba-c) and NF-L, (green colour in Ba, b and d) in the nerve fibres, mesenchyme (m) and enamel organ of an E14.5 tooth germ. Co-localization of Nogo-A and NF-L is visualised by the yellow colour in the merged images (Ba and Bb). DAPI (blue colour) marks the nuclei of the cells. The asterisk in (Ba) indicates the area of higher magnification images in (Bb-d). **(C)** Immunohistochemistry showing the distribution of Nogo-A in the nerve fibres and odontoblasts (o) of a P25 mouse molar. **(D)** Immunofluorescent staining of Nogo-A (red colour in Da-c), NF-L (green colour in Da, b and d), and DAPI (blue colour) in a P25 WT mouse molar. Yellow colour shows the co-localization of the Nogo-A and NF-L proteins in the merged images (Da and b). The asterisk in (Da) indicates the area of higher magnification images in (Db-c). **(E-G)** Confocal immunofluorescent images of Nogo-A (red colour), peripherin (periph), parvalbumin (PV), and calcitonin gene-related peptide (CGRP) (green colour in E, F, G, respectively) showing their localisation in the nerve fibres of a P25 molar. Co-localisation of Nogo-A with any other protein in merged images (Ea and b, Fa and b, Ga and b) is seen in yellow. DAPI (blue colour) indicates the cell nuclei. The asterisks in (Ea, Fa, Ga) indicate the areas of higher magnification images in (Eb-d, Fb-d, Gb-d), respectively. Additional abbreviations: dp, dental papilla; Mc, Meckel cartilage; d, dentine; p, dental pulp; oe, oral epithelium. Scale bars: 100 µm in (A, Ba, C, Da, Ea, Fa, Ga), 50 µm in (Bb-d, Db-d, Eb-d, Fb-d, Gb-d).

In the dental pulp of the fully erupted teeth, a strong Nogo-A staining was detected in the nerve fibres (**Fig. 2C** ), identified by NF-L expression (**Fig. 2D** ). Nogo-A-positive fibres were also detected in the sub-odontoblastic neuronal plexus in close proximity to the odontoblasts (**Fig. 2Db-d** ). Since the innervation of the tooth is heterogeneous and composed of several sensory systems (70), we wished to determine which neuronal subtype expresses Nogo-A. For this purpose, we performed co-labelling in P25 WT lower molars with peripherin, parvalbumin (PV), and calcitonin gene-related peptide (CGRP), which constitute specific markers of the three sensory complexes (70) (**Fig. 2E-G** ). Peripherin was used to mark the mechano-nociceptive neurons, PV the mechano-sensitive (non-nociceptive) ones, and CGRP the second nociceptive neurons (70) of the dental pulp. Although Nogo-A labelled the majority of the fibres stained for peripherin (**Fig. 2E**) and PV (**Fig. 2F**), only a portion of CGRP-positive fibres were stained for Nogo-A (**Fig. 2G**), suggesting that the two nociceptive systems of the tooth differentially express Nogo-A. Therefore, in the pulp of postnatal teeth, Nogo-A is expressed by neurons of the mechano-nociceptive system (peripherin-positive / CGRP-negative) and of the mechano-sensitive system (PV-positive / CGRP-negative), while Nogo-A expression in CGRP-nociceptive neurons (CGRP-positive / PV-negative / peripherin-negative) is limited.

### Reduced neuronal complexity in the teeth of Nogo-A KO mice

Next, we wanted to explore how Nogo-A regulates the innervation network of the tooth. For that purpose, we analysed Nogo-A knockout mice (Nogo-A KO), in which the Nogo-A isoform of the *Rtn4* gene is deleted (54). Previously, these mice demonstrated branching defects in the neuronal network of the developing embryonic limbs (19). Dental pulps were isolated from WT and Nogo-A KO mice at different stages of development and processed for whole-mount fluorescent immunostaining for NF-L and peripherin. These stages include early stage of neuronal invasion in the dental pulp (P4), pre-eruption tooth stage in which the formation of neuronal network has started to be established (P7), and the erupted tooth stage (P25) (**Fig. 3** ). In the P4 WT molars (**Fig. 3A** ), the pioneer neuronal branches had already invaded the dental pulp and formed the main branch which reached the odontoblastic layer (**Fig. 3Aa, b** ). In contrast, in the P4 Nogo-A KO molars (**Fig. 3B** ), fewer neuronal branches invaded the dental pulp, and although the main neuronal branch reached the odontoblastic layer, it showed less ramification compared to WT (**Fig. 3Ba, b** ). In the P7 WT molars (**Fig. 3C** ), additional secondary neuronal branches had entered the dental pulp, and the network density had highly increased in the sub-odontoblastic region (**Fig. 3Ca, b** ). In the P7 Nogo-A KO molars (**Fig. 3D** ), the secondary branches were also present, but displayed reduced branching and sparser network in the sub-odontoblastic region compared to WT (**Fig. 3Da, b**). In the P25 WT molars (**Fig. 3E**), thick nerve bundles ran the whole length of the tooth roots and most of the tooth crown (**Fig. 3Ea-c** ). Nerves branched extensively in the pulp horns of the tooth crown (**Fig. 3Ea, b)** and to a lesser degree in the tooth root area (**Fig. 3Ea, c** ). In the P25 Nogo-A KO molars (**Fig. 3F** ), the neuronal network was less dense in the tooth crown (**Fig. 3Fa, b**), and the neuronal bundles in the tooth root area were less thick and dense (**Fig. 3Fa, c**).

**Figure 3:**
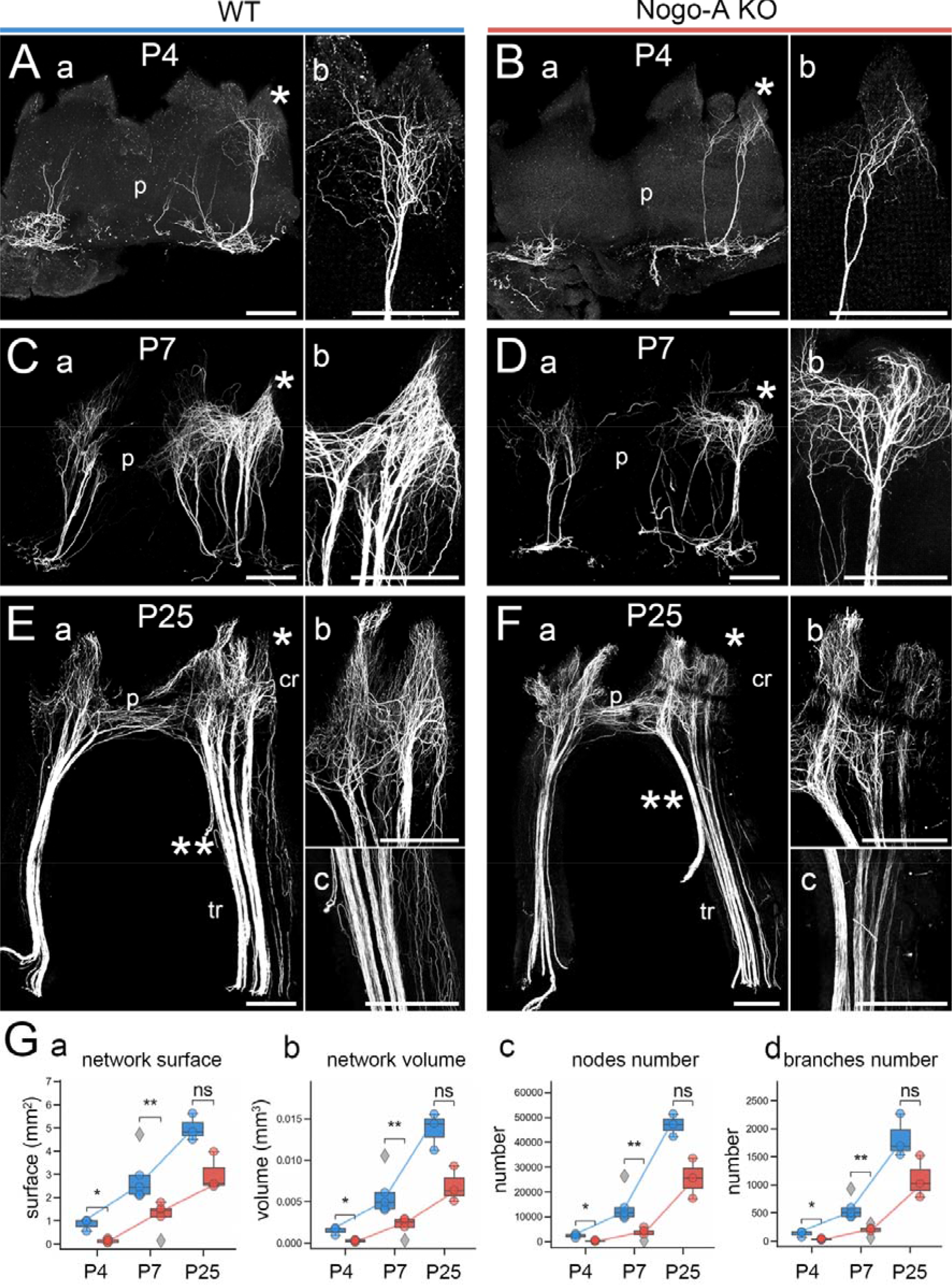
Effects of the Nogo-A deletion on tooth innervation. . **(A-F)** Representative whole-mount fluorescent staining images of combined NF-L and peripherin expression in the WT (A, C, E) and Nogo-A KO (B, D, F) first lower molars, at P4 (A, B), P7 (C, D), and P25 (E, F). The single asterisks in (Aa-Fa) indicate the areas in the tooth crown (cr) shown in higher magnification in (Ab-Fb), respectively. The double asterisks in (Ea, Fa) indicate the areas in the tooth root (tr) shown in higher magnification in (Ec, Fc), respectively. Notice the reduction of the neuronal branching and sparser network in the dental pulp of Nogo-A KO. **(G)** Quantification of the network surface (Ga), volume (Gb), number of nodes (Gc), and number of branches (Gd) in WT (blue colour) and Nogo-A KO (red colour) teeth at P4, P7, and P25. Statistical analysis: Two-sided Mann–Whitney U test (ns, p > 0.05; *, p ≤ 0.05; **, p ≤ 0.01; ***, p ≤ 0.001). Additional abbreviation: p, dental pulp. Scale bars: 200 µm.

Thereafter, we quantified the surface, volume, and number of nodes and branches in all these stages in both WT and Nogo-A KO teeth (**Fig. 3G** ). Although the data exhibited high biological variability, 75% of the data set showed a similar trend. At all stages analysed, the innervation network of the Nogo-A KO molars was reduced, both in terms of surface (**Fig. 3Ga**), volume (**Fig. 3Gb** ), and number of nodes (**Fig. 3Gc** ) and branches (**Fig. 3Gd** ). The reduced complexity of the innervation network within the dental pulp of Nogo-A KO teeth suggests a delay in maturation compared to the innervation of WT teeth. Indeed, the neuronal network in P7 Nogo-A KO teeth is comparable to that observed in P4 WT teeth. Similarly, both the neurite network density and complexity in the dental pulp of P25 Nogo-A KO teeth are still lower when compared to the P25 WT teeth, thus suggesting either a long lasting or a permanent neuronal deficit created by the neuronal maturation delay.

### Genetic and/or pharmacological Nogo-A inhibition affects the outgrowth of tgg-derived neurites and their interaction with the tooth germs in vitro

Nogo-A was shown to affect the outgrowth of *in vitro* cultured dorsal root ganglia (DRG), the tgg counterpart responsible for the somatosensory innervation of the body (19). Therefore, we analysed the effects of Nogo-A on the outgrowth of embryonic tgg explants obtained from WT and Nogo-A KO mice *in vitro* (**Fig. 4**). After two days in culture, WT tgg projected a dense network of axon processes accompanied by migration of cells from the tgg (**Fig. 4A**). Fluorescent double immunostaining showed that although many of these neuronal projections co-expressed Nogo-A and NF-L (**Fig. 4Ac-e, F, G**), a significant number of neurons were marked by exclusive Nogo-A or NF-L labelling (**Fig. 4Ac, Fa, Ga** ). Furthermore, the intensity of the Nogo-A labelling varied along the length of many neuronal processes, contrasting to the mainly uniform NF-L labelling observed in these samples (**Fig. 4Ac, Fa, Ga**). Tgg from Nogo-A KO mouse embryos produced a more extensive network of axonal outgrowths (**Fig. 4Ba-c, e** ) when compared to WT tgg (**Fig. 4Aa-c, e** ). For quantification, we segmented the axonal outgrowths from the bright field images (**Fig. 4C-D**) and performed a Sholl analysis (**Fig. 4E** ), which confirmed that the Nogo-A KO tgg grew more and longer neurites compared to the WT tgg. In addition, the cell bodies of neurons migrating out of Nogo-A KO tgg during the two days of culture were found further away from the tgg (**Fig. 4Bb, c**) when compared to neuronal cell bodies of WT tgg (**Fig. 4Ab, c**).

**Figure 4:**
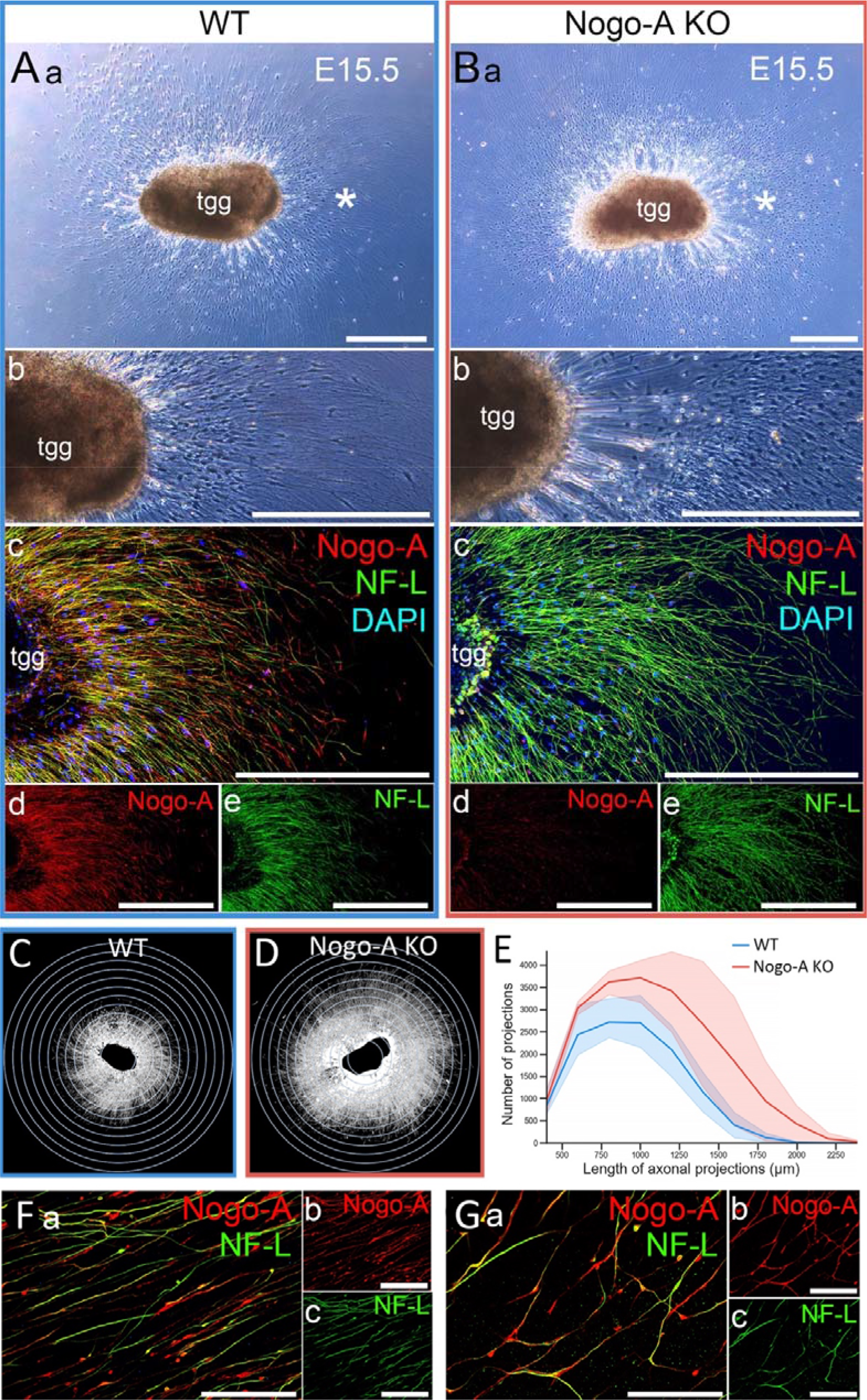
Neurite outgrowths from *in vitro* cultured tgg explants of WT and Nogo-A KO mouse embryos. *In vitro* culture of tgg explants obtained from WT (A, C, F, G) and Nogo-A KO (B, D) mouse embryos at E15.5. **(Aa, b and Ba, b)** Bright field images of the WT (Aa, b) and Nogo-A KO (Ba, b) trigeminal ganglia after 2 days in culture. The asterisks in (Aa and Ba) indicate the areas shown in higher magnification in (Ab-e and Bb-e), respectively. Note the higher extension of the neuronal projections from tgg derived from the Nogo-A KO mouse embryos. Also note the presence of neuronal cell bodies further away from the Nogo-A KO tgg body (compare Ab to Bb). **(Ac-e, and Bc-e)** Immunofluorescent staining of Nogo-A (red colour) and NF-L (green colour) in the cultured WT (Ac-e) and Nogo-A KO (Bc-e) tgg. Yellow colour shows the co-localisation of Nogo-A and NF-L. DAPI (blue colour) marks the nuclei of the cells (Ac, Bc). Note the longer neuronal projections of the NF-L-positive neurons originated from the Nogo-A KO tgg (compare Ae to Be). **(C-D)** Segmentation of the developing neuronal network from the cultured WT (C) and Nogo-A KO (D) tgg explants in (Aa) and (Ba), respectively. **(E)** Sholl analysis of the segmented network from (C) (blue colour) and (D) (red colour). **(F and G)** Immunofluorescent staining of Nogo-A (red colour) and NF-L (green colour) in the neuronal network of the cultured WT tgg. Co-staining of Nogo-A and NF-L is seen in yellow. Note that the Nogo-A-positive neurons are not always co-stained with NF-L. Abbreviation: tgg, trigeminal ganglion. Scale bars: 500 µm in (Aa-e and Ba-e), 50 µm in (Fa-c and Ga-c).

We then analysed the *in vitro* effects of Nogo-A on tooth innervation in co-cultures of tgg and tooth germ explants (**Fig. 5**). Embryonic tooth germs strongly repel neurite outgrowth from tgg in co-culture (71). To identify and quantify the neurite outgrowth from tgg, we dissected tgg from R26-mTmG embryos that ubiquitously express the red fluorescent protein (RFP) and analysed the RFP-positive tgg neurite outgrowth. In a first set of experiments (**Fig. 5A-B**), embryonic WT molar tooth germs were placed approximately 1.5 mm away from the tgg (**Fig. 5Aa**) and treated with either an IgG-control antibody (**Fig. 5Ab**) or an anti-Nogo-A antibody (**Fig. 5Ac**). The tgg neurite outgrowth was segmented using the RFP signal (**Fig. 5Ad, e** ) and its branching complexity was quantified by a Sholl analysis (**Fig. 5Af**). The control-IgG treated tgg generated on average less than 20 projections that were long enough to reach the tooth germ (**Fig. 5Af**). However, none of these were projecting in the direction of the tooth germ (**Fig. 5Ab, d** ). Tgg treated with the anti-Nogo-A antibody projected on average 10 times more and 2 times longer axons than the control-treated tgg (**Fig. 5Af** ). Several of these projections reached and contacted the tooth germ in these cultures. They created a dense network around the tooth germ that was in part directly contacting the surface of the tooth germ (**Fig. 5B**). While the axons did not appear to enter the dental mesenchyme, the continuous inhibition of Nogo-A did promote the formation of an extensive and complex innervation network around the embryonic tooth germ. Similar results were observed when the tgg was cultured together with both WT and Nogo-A KO tooth germs in a microfluidic device (**Fig. 5Ca-c**). In this setup, the tooth germ of WT mouse embryos repealed the neurites originating from the tgg of WT mouse embryos (**Fig. 5Cb**). In contrast, the WT tgg outgrowths were clearly directed towards the tooth germ of Nogo-A KO mouse embryos (**Fig. 5Cc**). However, the overall tgg outgrowth pattern was unaffected in terms of length and thickness of the fibres by the absence of Nogo-A in the tooth germs. Finally, in another set of experiments, we cultured tooth germs from either WT **(Fig. 5Da-c**) or Nogo-A KO **(Fig. 5Dd-f**) mouse embryos in immediate proximity to the R26-mTmG tgg. In this setup, we observed that the Nogo-A KO tooth germs permitted close contact with many tgg projections compared to WT tooth germs that showed very little interaction with the tgg projections (**Fig. 5Dg**).

**Figure 5:**
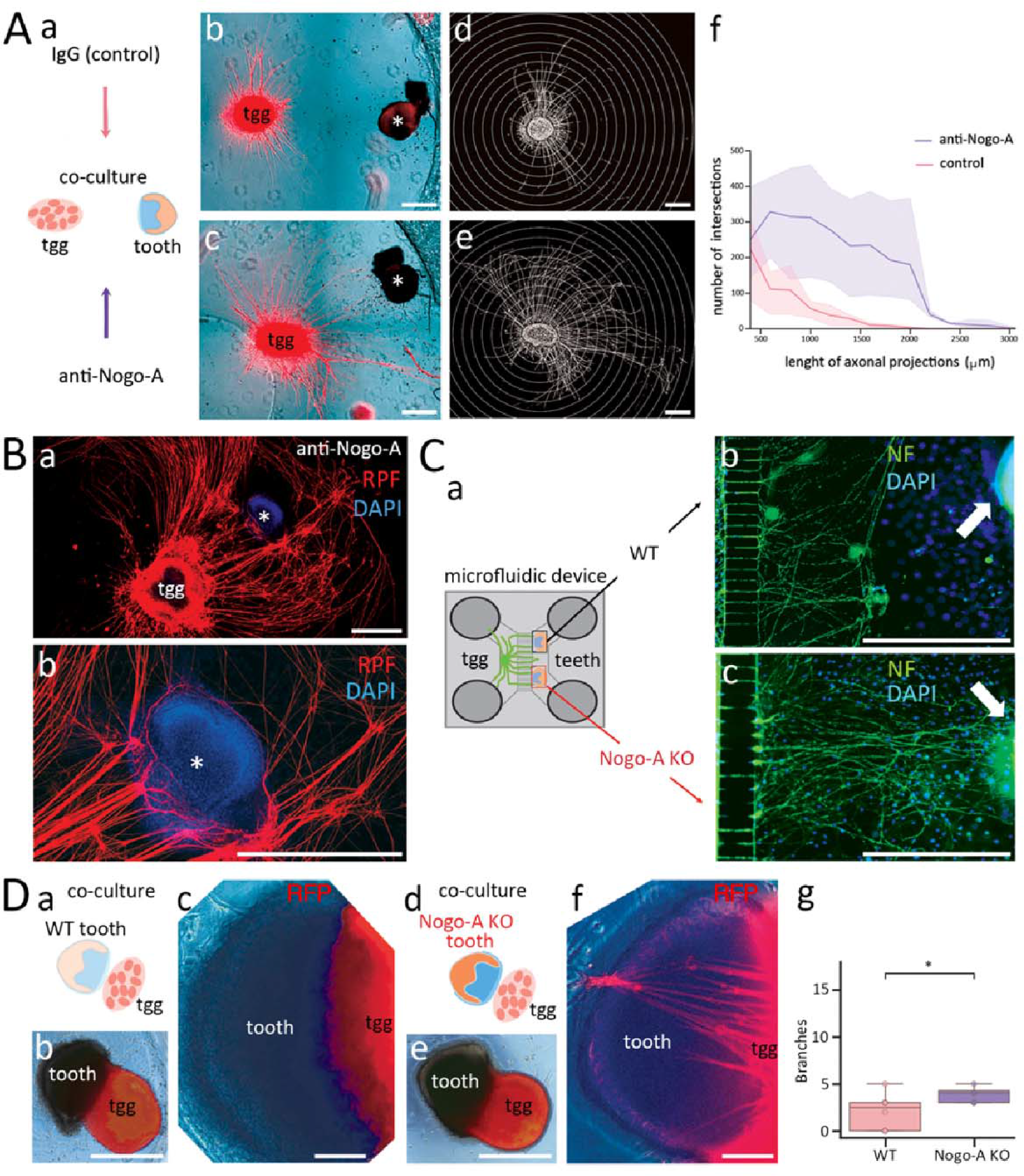
Nogo-A affects the interaction between tooth germs and tgg neurons. . **(Aa)** Schematic representation of co-cultures of tgg from a R26-mTmG mouse embryos and tooth germs from WT mouse embryos. The co-cultured explants were treated with either IgG (control) or anti-Nogo-A antibodies for 6 days. **(Ab,c)** Representative images of tgg and tooth germs co-cultures treated with IgG (Ab), and anti-Nogo-A antibodies (Ac). The asterisks indicate the tooth germs. Note the repulsion of neurons from the tooth germ in the control culture conditions (Ab) and the opposite effect created upon treatment with Nogo-A antibodies (Ac). **(Ad,e)** Segmentation of the neuronal network following 6 days of culture based on the RFP channel images. Note the obvious repulsion of neurons from the tooth germ in the control culture conditions (Ad), and the evident attraction of neurons towards the tooth germ after treatment with Nogo-A antibodies (Ae). **(Af)** Sholl analysis of the segmented network from (Ad) (red colour) and (Ae) (purple colour). **(Ba)** RFP staining (red colour) of a co-cultured tgg with a tooth germ in presence of anti-Nogo-A antibodies. DAPI (blue colour) stains the nuclei of the cells. The asterisk indicates the tooth germ. **(Bb)** Higher magnification of (Ba) showing the dense neuronal network surrounding the tooth germ. **(Ca)** Schematic representation of a microfluidic device allowing the co-culture of tgg from WT mouse embryos and tooth germs from WT and Nogo-A KO mouse embryos. **(Cb)** Neurofilament (NF) immunofluorescent staining (green colour) showing a directional growth of the tgg-derived neurons far away from the WT tooth germ (arrow) after 10 days of culture. DAPI (blue colour) stains the nuclei of the cells. **(Cc)** NF immunofluorescence (green colour) showing a growth of the tgg-derived neurons towards the Nogo-A KO tooth germ (arrow). **(Da)** Schematic representation of the tgg and tooth germs from WT mouse embryos co-cultures. **(Db)** Four-day co-cultures of tgg (red colour) isolated from R26-mTmG mouse embryos and tooth germs isolated from WT mouse embryos. **(Dc)** Higher magnifications of (Db) showing the absence of neuronal projections from the tgg towards the tooth germ. **(Dd)** Schematic representation of the tgg and tooth germs from Nogo-A KO mouse embryos co-cultures. **(De)** Four-day co-cultures of R26-mTmG tgg (red colour) and Nogo-A KO embryonic tooth germs. **(Df)** Higher magnifications of (De) showing numerous neuronal projections from the tgg towards the tooth germ. **(Dg)** Quantification of the number of neuronal projections formed from the tgg towards the immediately adjacent WT (red colour) and Nogo-A KO (purple colour) tooth germs. Statistical analysis performed by the Two-sided Mann–Whitney U test. ns, p > 0.05; *, p ≤ 0.05; **, p ≤ 0.01; ***, p ≤ 0.001. Scale bars: 500 µm in (A, B, C, Db, De), 100 µm in (Dc, Df).

### Gene expression changes in the tgg of Nogo-A KO embryonic and postnatal mice

In the last set of experiments, we investigated the gene expression changes in the E14 and P25 tgg of Nogo-A KO mice by extraction of total RNA and subsequent bulk RNA sequencing (**Fig. 6**). The data confirmed the deletion of the Nogo-A isoform in the Nogo-A KO mice, as no mRNA expression of the Nogo-A isoform was detected in Nogo-A KO tgg at either stage (**Fig. 6A** ). However, Nogo-A deletion affected the expression of the other *Rtn4* isoforms, especially in the postnatal stage where both Nogo-B and Nogo-C isoforms were upregulated in Nogo-A KO tgg (**Fig. 6A**).

**Figure 6:**
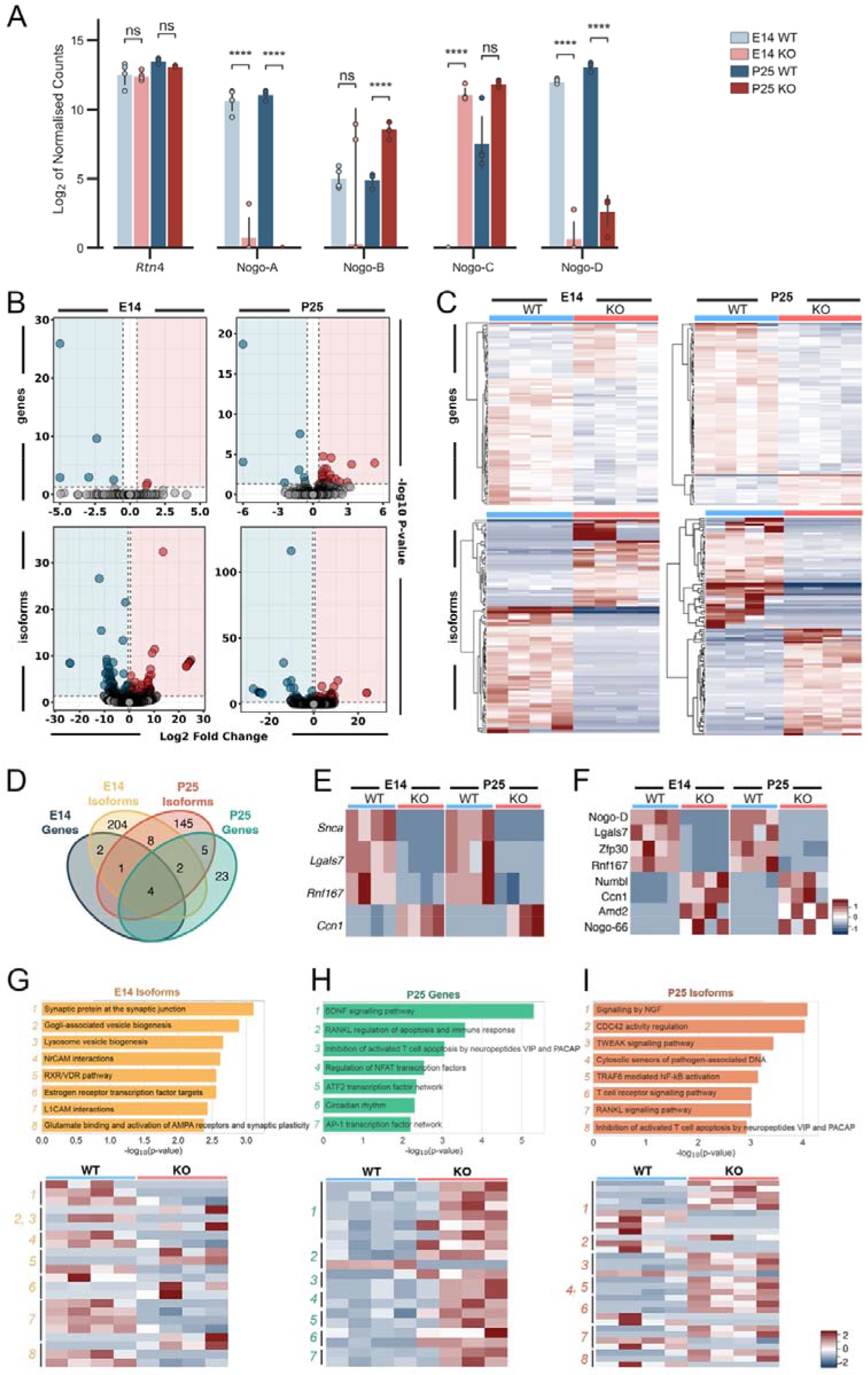
Bulk RNA sequencing analysis in tgg of WT and Nogo-A KO embryonic and postnatal mice. **(A)** Expression analysis of the *Rtn4* gene and its main isoforms Nogo-A, Nogo-B, Nogo-C, and Nogo-D in the tgg of WT (blue colour) and Nogo-A KO (red colour) mice at E14 (light colour) and P25 (dark colour) stages. Expression levels are shown as the log_2_ of gene- and isoform-specific counts normalised to the total count of each sample. **(B)** Volcano plots of the significantly downregulated (p < 0.05 and log_2_(ratio) < -0.5) and upregulated (p < 0.05 and log_2_(ratio) > 0.5) genes (upper row) and isoforms (bottom row) in the E14 (left panel) and P25 (right panel) Nogo-A KO tgg. **(C)** Heatmap of the top 100 differentially expressed genes (upper row) and isoforms (bottom row) at E14 (left panel) and P25 (right panel). **(D)** Venn diagram showing the distribution of differentially expressed genes and isoforms in E14 and P25 Nogo-A KO tgg. **(E)** Heatmap showing changes in the expression of the differentially expressed genes shared between the E14 and P25 Nogo-A KO tgg. **(F)** Heatmap showing changes in the expression of the differentially expressed isoforms shared between E14 and P25 Nogo-A KO tgg. **(G-I)** Enrichr pathway analysis (upper row) and heatmaps of the genes of interest (bottom row) using the NCATS BioPlanet 2019 database of the differentially expressed isoforms at E14 (G), the differentially expressed genes at P25 (H), and the differentially expressed isoforms at P25 (I).

Analysis of E14.5 and P25 Nogo-A KO tgg showed moderate changes in gene expression levels when compared to WT control (**Fig. 6B** ). At E14.5, *Egr1* and *Ccn1*, which encode proteins affecting neuroblast proliferation and modulate their methylome (72, 73) were significantly upregulated (**Suppl. Table 1** ). Five genes, including *Snca, Mmrn1, Lgals7* and *Rnf167,* were significantly downregulated at the same stage (**Suppl. Table 2**). In tgg of P25 Nogo-A KO mice, genes associated with neurotrophin downstream signalling (*Btg2*, *Egr2*, *Fosb*, *Junb*, *Sgk1*), RNA processing (*Prpf39*, *Zcchc7*), circadian clock (*Per1*, *Per3*, *Ryr1*), neuronal functions (*Cap89*, *Rassf10),* cell cycle (*Cep290*, *Mki67),* extracellular matrix (*Cdh19*, *Lpp*, *Ccn1*, *Col24a1*, *Fcgbp)* and transcriptional regulation (*Hmgb2*, *Csrnp1*, *Nfkbia)* were significantly upregulated (**Suppl. Table 3** ). Interestingly, seven genes that were downregulated at this stage are also involved with neuronal functions (*Txnip*, *Zfp36*), cell cycle (*Rnf167*), extracellular matrix (*Lgals7*), and transcriptional regulation (*Ier5*) (**Suppl. Table 4** ). Differential expression analysis of gene isoforms in our samples enables deeper insight into molecular differences caused by Nogo-A deletion. Indeed, these analyses demonstrated broader effect of Nogo-A deletion on the molecular profile of tgg when compared to differential expression analyses of individual genes (**Fig. 6C-D** and **Suppl. Table 5-8**).

Next, we wanted to identify which impacted genes are shared between the two developmental stages (**Fig. 6D** ). These genes include *Snca*, *Lgals7,* and *Rnf167* which were downregulated, and *Ccn1* which was upregulated (**Fig. 6E** ). The *Lgals7* gene encodes galectin, a protein that is secreted in the extracellular matrix, and promotes the proliferative, migratory, and wound healing properties of periodontal fibroblasts (77, 78). Rnf167 is an ubiquitin ligase which acts as modulator of neurotransmission, endosomal trafficking, and lysosome positioning (79). Ccn1, a matricellular protein, is a known downstream target of RhoA, which participates in cell proliferation, migration, and differentiation, depending on the cellular context (80–82). Interestingly, Ccn1 and Nogo-A share several interaction partners such as TrkA, integrins, and heparan sulphate proteoglycans (HSPGs) (81, 83–85). *Snca* was significantly downregulated in the Nogo-A KO at both stages, but this is likely due to a genomic deletion on chromosome 6 that includes the *Snca* and *Mmrn1* (downregulated at E14.5 only) loci and is present in the Nogo-A KO mouse strain (86). Additionally, the Nogo-D isoform and the isoforms of *Zfp30* and *Rnf167* were downregulated at both stages, while the short Nogo-66 fragment and the isoforms of *Numbl*, *Ccn1* and *Amd2* genes were upregulated (**Fig. 6F**).

Enrichment analyses demonstrated that genes and isoforms whose expression is altered at E14 are related to dysregulation of synaptic function (**Fig. 6G** and **Suppl. Table 9**), either by encoding synaptic proteins, or proteins involved in vesicle biogenesis, and interactors of NrCAM, which is involved in synapse formation and axon growth and guidance (74). Greater number of genes and isoforms affected in Nogo-A KO tgg at P25 belonged to the signalling network of neurotrophins BDNF and NGF (**Fig. 6H-I** and **Suppl. Tables 10-11**). Regulators of the activity of CDC42, that belongs to the Rho GTPase family which regulates the neurite growth and plasticity (75), were also dysregulated in the Nogo-A KO tgg (**Fig. 6I** and **Suppl. Table 11**). Notably, in contrast to the compensation mechanisms that have been reported in the CNS of adult Nogo-A KO mice, we found no upregulation of other myelin-associated inhibitors or axon guidance molecules in the Nogo-A KO tgg, such as semaphorins, ephrins, and MAG (76).

## Discussion

Nogo-A is a well-known inhibitor of neurite outgrowth and plasticity in the central nervous system (CNS) (14, 15). It is also involved in the regulation of non-neuronal biological processes such as angiogenesis and vascular remodelling, macrophage activation, and stem cell behaviour (87). Surprisingly, the functions of Nogo-A in the highly innervated dental tissues were not assessed until recently (88). In the present study, we analysed the effects of Nogo-A deletion in the embryonic and postnatal trigeminal ganglia (tgg), and for the first time we assessed its roles in tooth innervation. During embryogenesis, Nogo-A is expressed in the tgg and tgg-derived neurites that surround, but do not enter, the connective tissues of the developing tooth germs. During postnatal development, the trigeminal axons innervate the connective tissues of the tooth while they continue to express Nogo-A. Within the dental pulp of young adult mice, Nogo-A-expressing neurites are part of the sub- odontoblastic neuronal plexus, in close association with the dentine-producing odontoblasts. These results are in agreement with our previous findings showing Nogo-A expression in the neurons of human permanent teeth (88).

Tooth innervation is regulated by neuroregulatory molecules that either attract or repulse the growing trigeminal nerves (43). Nerve growth factor (NGF) and glial-derived neurotrophic factor (GDNF) are the most extensively studied neuro-attractant molecules during tooth innervation (43). Increased NGF expression in the developing tooth germs leads to the attraction of growing trigeminal neurons into the dental pulp tissue (46, 89). Similarly, increased GDNF expression in the postnatal dental mesenchyme is responsible for the guidance of neurons towards the sub-odontoblastic area (89–91). It has been shown that the developing trigeminal axons express p75^NTR^ and TrkA, which are receptors for NGF, as well as the GDNF-receptor Ret (92). Previous studies have demonstrated that NGF or GDNF deletion drastically decreases tooth innervation (48, 93, 94). In contrast to these two neuro-attractant molecules, semaphorin 3A (Sema3A) has a repulsive effect on tooth innervation (50, 51). Sema3A is highly expressed in the embryonic dental mesenchyme, where it inhibits the growth of trigeminal axons that express its receptors (50). A decrease in Sema3A expression in the postnatal dental pulp tissue is associated with the initiation of its innervation by trigeminal neurons. Its functional relevance was further demonstrated by the observation of ectopic and premature innervation of developing tooth germs in Sema3A KO mice (95). Taken together, these studies show that a balance between neuro-attractive and neuro-repulsive molecules is needed to regulate proper tooth innervation. Nogo-A is an integral part of this process, as its actions are strongly intertwined with those of semaphorins and neurotrophins. p75^NTR^ can act as a receptor for both Nogo-A and NGF (96), and most of the above-mentioned molecules exert their action on axonal growth by modulating the activation of the ROCK pathway. Sema3A and Nogo-A both mediate the disorganisation of the cytoskeleton via RhoA and ROCK activation (10, 97), whereas NGF has the opposite effect and inactivates RhoA and ROCK (97). Our present findings demonstrate the importance of Nogo-A in tooth innervation. Indeed, Nogo-A deletion strongly reduced the volume of the neuronal network and the number of neuronal nodes within the dental pulp tissue. This results in the formation of a simple, less complex neuronal network in the teeth of adult Nogo-A KO mice. Similar neuronal patterning defects have already been reported for other tissues and organs of Nogo-A KO mice, such as the limbs (19). Nogo-A on the surface of the growing fibres of the tgg as well as Nogo-A present in the tooth tissue may influence bundling and branching as well as guidance and growth speed along fibre bundles. An intrinsic neuronal effect is suggested by the observation that neurites emerging from tgg cultured in the presence of blocking anti-Nogo-A antibodies were thicker and longer. A similar effect has been previously observed in DRGs cultured with Nogo-A blockers

(19). On the other hand, the inhibition of Nogo-A activity also drastically affected the interactions between the tgg-derived neurites and the developing tooth germs, which are known to repel trigeminal neurites (71). Although tgg-derived neurites do not project towards the tooth germs in normal conditions, these same neurites were attracted towards the dental explants and formed a dense network around the germs upon treatment with anti-Nogo-A antibodies. Similarly, tgg-derived neurons from wild-type mice emitted more projections towards the tooth germs of Nogo-A KO mice compared to the WT tooth germs. Identifying molecules involved in the establishment of connections between the peripheral nerves and dental tissues is essential for the development of novel therapeutic approaches for optimal recovery of tooth physiology. Indeed, in teeth like in many organs, the innervation is emerging as a key player in organ development, homeostasis, repair and/or regeneration (88–91). Our *in vitro* results suggest the potential use of Nogo-A blockers for the treatment of dental tissues in a clinical setting. This statement is reinforced by our recent finding showing that Nogo-A in involved in the differentiation process of human dental pulp stem cells towards the osteogenic, neurogenic, and adipogenic phenotypes (88). Therefore, our previous and present results encourage the development of therapeutics targeting Nogo-A for dental clinical treatments.

The RNA sequencing analysis in tgg showed that Nogo-A deletion moderately affects the tgg transcriptome. This contrasts with what was previously reported in the adult CNS, where the expression of other repulsive axon guidance molecules was upregulated in compensation for Nogo-A deletion (76). The absence of such compensation in the tgg of Nogo-A KO animals suggests different functions for the Nogo-A molecule in the cranial nerves and the CNS. Nevertheless, the enrichment analyses revealed defects in synaptic function in the embryonic Nogo-A tgg, with the potential dysregulation of synaptic proteins and Golgi- and lysosome vesicle biogenesis (103). This is further supported by observed impact on the expression of interactors of NrCAM, a neuronal cell adhesion molecule involved in synapse formation and axon growth and guidance (74). In the postnatal Nogo-A tgg, signalling components of NGF and BDNF were affected, as well as the CDC42, a sibling of RhoA in the Rho GTPase family. Members of the Rho GTPase family act as regulators of morphological neuroplasticity and are downstream signal transducers for several repulsive guidance molecules including Nogo-A (9, 10, 75). Together, these results suggest that Nogo-A regulates synaptic function and axon growth and guidance in the embryonic tgg and tooth and interferes with neurotrophin and Rho signalling in the postnatal tgg.

In conclusion, the present results reveal for the first time the crucial role of Nogo-A in regulating tooth innervation (**Fig. 7**) and indicate its potential for future clinical applications for teeth and other peripheral innervated organs.

**Figure 7:**
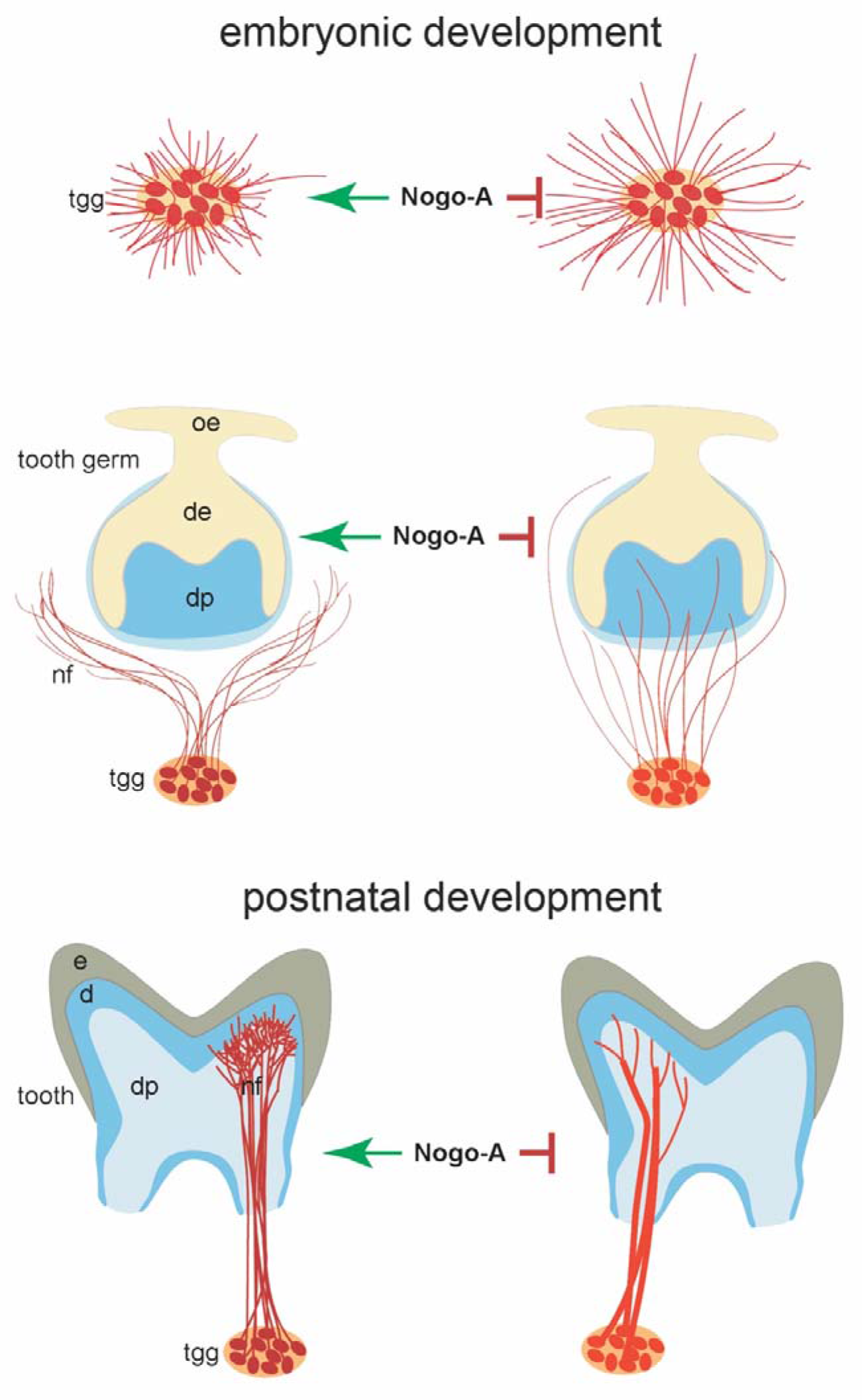
Schematic representation summarizing the functional roles of Nogo-A in trigeminal ganglion and tooth innervation. Absence of Nogo-A during the embryonic development contributes to longer neuronal projections from the trigeminal ganglion and their attraction towards the developing tooth germs. At postnatal stages, absence of Nogo- A is responsible for the increased thickening of the neurons, their reduced neuronal branching and sparser network in the dental pulp. Abbreviations: d, dentine; de, dental epithelium; dp, dental pulp; e, enamel; nf, nerve fibres; tgg, trigeminal ganglion.

## Acknowledgements

Funds: This project was financed by the SNSF (grant number: 31003A_179389) and by institutional funds from the University of Zurich (UZH). We are grateful to Argyro Lamprou and Maria Lydia Ioannides for helping with the histological sections, and Dr Isaac Bugueno for helping with the draft writing.

## Contributions

Conceptualization: T.A.M., M.E.S; methodology: T.A.M., L.P., An.B., C.F.L, P.P., M.E.S.; data analysis: T.A.M., M.E.S, L.P., An.B., I.D.S., Al.B., C.F.L.; validation: T.A.M., L.P., An.B., M.E.S.; formal analysis: T.A.M., L.P., An.B., I.D.S., Al.B.; investigation: L.P., C.F.L., P.P.; resources, T.A.M., M.E.S.; data curation: T.A.M., L.P., An.B., P.P., M.E.S.; writing draft: T.A.M., L.P., An.B.; draft editing: T.A.M., L.P., An.B., P.P., M.E.S; reading and validation of final draft: T.A.M., M.E.S, L.P., An.B., I.D.S., Al.B., C.F.L., P.P.; visualization: T.A.M., L.P., An.B.; supervision: T.A.M., An.B., P.P.; project administration: T.A.M.; funding acquisition, T.A.M. All authors have read and agreed to the published version of the manuscript.

